# Identification and Diagnostic Potential of Pyroptosis-Related Genes in Endometriosis: A Novel Bioinformatics Analysis

**DOI:** 10.1101/2024.09.23.614461

**Authors:** Piaopiao Teng, Li Wang, Caiyun Ding, Kaili Gu, Xianchen Luo, Chang Su, Guantai Ni, Yuanyuan Lyu, Jin Ding

**Affiliations:** Department of Obstetrics and Gynecology, First Affiliated Hospital of Wannan Medical College, 241000Wuhu, Anhui, P.R. China.; Modern Health and Wellness Industry College, Anhui Sanlian University, No. 47 He’an Road, 230601Hefei, Anhui, P.R.China; School of Basic Medicine, Wannan Medical College, 241002 Wuhu, Anhui, P.R.China

**Author notes:** Corresponding Author (Yuanyuan Lyu), (Jin Ding). These authors contributed equally (Piaopiao Teng, Li Wang).

**Keywords:** Endometriosis, Pyroptosis-Related Genes, Diagnostic models, immune cell infiltration, Bioinformatics

## Abstract

**Objective:** This study aimed to identify and analyze potential signatures of pyroptosis-related genes in EMs.

**Methods:** Transcriptomic datasets related to endometriosis were retrieved from the GEO databases (GSE7305, GSE7307, and GSE11691). Differential gene expression analysis was performed to identify pyroptosis-related differentially expressed genes (PRDEGs) by intersecting DEGs with a curated list of PRGs. Various bioinformatics tools were employed to explore the biological functions and pathways associated with PRDEGs.

**Results:** We identified 26 PRDEGs from combined datasets and constructed an EMs diagnostic model using LASSO regression based on pyroptosis scores. The model included 5 DEGs: KIF13B, BAG6, MYO5A, HEATR, and AK055981. Additionally, 21 Key Module Genes (KMGs) were identified, leading to the classification of 3 distinct EMs subtypes. These subtypes were analyzed for immune cell infiltration, revealing a complex immune landscape in EMs.

**Conclusions:** This study reveals pyroptosis’ crucial role in EMs and offers a novel diagnostic model based on pyroptosis-related genes. Modulating pyroptosis may provide a new therapeutic approach for managing EMs.

## 1. Background

Endometriosis is defined by the presence of endometrium-like epithelium and/or stroma outside the uterus(Becker et al., 2022). It reduces the quality of life about 10% of women in childbearing age and results in symptoms such as pain and infertility(Wang et al., 2020). Endometriosis is characterized as an inflammatory condition, wherein the presence of ectopic lesions can precipitate pelvic inflammation, thereby facilitating the further development of these ectopic endometrial lesions. The recurrent inflammatory responses associated with this condition result in an abnormal elevation of inflammatory cytokines, which are essential for the attachment, growth, specialization, and invasion of endometrial lesions(Zhang et al., 2018). However, the fundamental mechanisms underlying endometriosis remain poorly understood, and further research is required to elucidate its pathogenesis.

Pyroptosis is an inflammatory form of programmed cell death initiated by a variety of disease-causing factors. It has received increasing attention because of the association with innate immunity and disease(Chen et al., 2023; Yu et al., 2021). It has been established that pyroptosis is induced within the stromal compartments of endometriotic lesions by bufalin. This induction is associated with increased activity of IL-1β and caspase-1 in stromal cells.(Cho et al., 2018). The release of prostaglandin E2 (PGE2) as a consequence of pyroptosis has been implicated in the advancement of endometriosis lesions by influencing cellular migration(Huang et al., 2022). Additionally, TRIM24 may play a role in the progression of endometriosis through the NLRP3/caspase-1/interleukin-1 (IL-1) mediated pyroptotic pathway, potentially via the ubiquitination of NLRP3(Hang et al., 2021). The findings of these studies indicate that pyroptosis plays a role in the pathogenesis of endometriosis.

The growing prevalence and utilization of gene chip sequencing technology have rendered microarray analysis a valuable and innovative approach for the identification of genes associated with susceptibility to endometriosis. Pyroptosis-related genes in endometriosis patients were sought to be identified through bioinformatics analysis based on transcriptome sequencing data in this study, thereby offering new insights for the diagnosis and treatment of the endometriosis.

## 2. Methods

### 2.1 Data collection

Endometriosis-related transcriptomic datasets GSE7305, GSE7307, and GSE11691, along with the validation dataset GSE25628, were retrieved from the Gene Expression Omnibus (GEO) database using the GEOquery package in R. All datasets comprised Homo sapiens endometrial samples. Batch effects across datasets were addressed using the surrogate variable analysis (sva) package in R, resulting in a combined dataset of 37 endometriosis and 42 normal samples.

A comprehensive list of pyroptosis-related genes was compiled from the GeneCards database and the Gene Set Enrichment Analysis (GSEA) database. Following the merging and removal of duplicate entries, a final set of 338 unique pyroptosis-related genes was established (Table S1).

### 2.2 Analysis of PRDEGs

Differential gene expression analysis between Endometriosis and Normal groups was performed using the DESeq2 package in R. Genes with an absolute log2 fold change (|logFC|) greater than 1 and an adjusted p-value (adj.p) less than 0.05 were considered differentially expressed (DEGs). The intersection of DEGs and the curated list of pyroptosis-related genes was identified and visualized using a Venn diagram to obtain PRDEGs.The results of the differential gene expression analysis were visualized using various R packages: ggplot2 for volcano plots, pheatmap for heatmaps, RCircos for chromosome localization, and pheatmap for correlation heatmaps (Chen et al., 2023). Correlation scatter plots were generated using ggplot2. Gene Ontology (GO) and Kyoto Encyclopedia of Genes and Genomes (KEGG) pathway enrichment analyses of PRDEGs were conducted using the clusterProfiler package in R.

### 2.3 Construction and Validation of a Diagnostic Model for Endometriosis

A pyroptosis score for each endometriosis patient was calculated using the single-sample Gene Set Enrichment Analysis (ssGSEA) algorithm. Patients were then stratified into high and low pyroptosis expression groups based on the median pyroptosis score. A diagnostic model for endometriosis was developed using the randomForest package in R. Subsequently, Least Absolute Shrinkage and Selection Operator (LASSO) regression analysis was performed using the glmnet package with the parameters set.seed(500) and family = “cox” to identify key features associated with endometriosis diagnosis. A nomogram was generated for visualizing the diagnostic model using the rms package. Finally, the diagnostic accuracy and discriminative power of the model were evaluated in the independent validation dataset GSE25628 by constructing decision curve analysis (DCA) plots using the ggDCA package.

### 2.4 GSEA(Gene Set Enrichment Analysis)

In this research, the genes in the Endometriosis group were initially categorized into two groups— those exhibiting high phenotypic correlation and those demonstrating low phenotypic correlation — according to their phenotypic correlation rankings. And then all genes in the Endometriosis group of the Combined Datasets were analyzed for GSEA by the R package clusterProfiler.

### 2.5 GSVA(Gene Set Variation Analysis)

The hallmark gene set (“h.all.v7.4.symbols.gmt”) was obtained from the Molecular Signatures Database (MSigDB). GSVA was performed on all genes within the endometriosis samples of the combined dataset to assess the enrichment of hallmark pathways. Differential pathway enrichment between the high and low pyroptosis expression groups was determined, with a significance threshold of p < 0.05.

### 2.6 WGCNA(Weighted correlation network analysis)

WGCNA was performed using the dedicated R package WGCNA to identify modules of highly correlated genes. The TOP2 modules exhibiting the strongest correlation (highest absolute r value) with pyroptosis expression were selected for further analysis. Genes within these modules were intersected with the DEGs identified in the high and low pyroptosis expression groups, respectively. Venn diagrams were generated to visualize the overlap between module genes and DEGs. The resulting intersecting genes from each module were designated as key module genes (KMGs).

### 2.7 PPI Network

A protein-protein interaction (PPI) network for the differentially expressed KMGs was constructed using the STRING database. A minimum required interaction score of 0.150 (low confidence) was applied to filter interactions. Subsequently, three algorithms within the cytoHubba plugin in Cytoscape were utilized to identify hub genes within the PPI network: Maximum Neighborhood Component (MNC), Degree, and Maximal Clique Centrality (MCC). The intersection of hub genes identified by each algorithm was determined and visualized using a Venn diagram. The resulting genes were designated as pyroptosis-related hub genes.

### 2.8 Construction of Regulatory Networks Involving Pyroptosis-Related Hub Genes

#### 2.8.1. ceRNA Network Construction

To explore the potential involvement of microRNAs (miRNAs) and circular RNAs (circRNAs) in regulating pyroptosis-related hub genes, miRNA-hub gene interactions were retrieved from the StarBase database. Subsequently, circRNA-miRNA interactions were also obtained from StarBase. The resulting mRNA-miRNA and miRNA-circRNA interactions were integrated to construct a competing endogenous RNA (ceRNA) network, which was visualized using Cytoscape software.

#### 2.8.2. Transcription Factor (TF)-mRNA Regulatory Network Construction

The ChIPBase database was utilized to identify TFs potentially regulating the pyroptosis-related hub genes. A TF-mRNA regulatory network was then constructed based on these predicted interactions and visualized using Cytoscape software.

#### 2.8.3. RNA-Binding Protein (RBP)-mRNA Regulatory Network Construction

The StarBase database was employed to predict RBPs targeting the pyroptosis-related hub genes. An mRNA-RBP regulatory network was subsequently constructed based on these predicted interactions and visualized using Cytoscape software.

#### 2.8.4. Drug-mRNA Regulatory Network Construction

The Comparative Toxicogenomics Database (CTD) was queried to identify direct and indirect drug targets associated with the pyroptosis-related hub genes, revealing potential interactions between these genes and various pharmacological agents. A drug-mRNA regulatory network was constructed based on these interactions and visualized using Cytoscape software.

### 2.9 Construction of disease subtypes of Endometriosis

Consensus Clustering (CC) is a resampling-based methodology employed to ascertain the membership of individual data points and their corresponding subgroup classifications, while also evaluating the validity of the identified clusters. The application of consensus clustering, utilizing the R package ConsensusClusterPlus, facilitates the identification of distinct disease subtypes of endometriosis, specifically in relation to the hub gene associated with pyroptosis.

### 2.10 Analysis of immune cell infiltration

The CIBERSORT algorithm was employed to estimate the relative abundance of immune cell subtypes within the endometriosis samples. Samples with a p-value less than 0.05 were selected for further analysis, and the immune cell infiltration matrix was generated using the LM22 eigengene matrix. The ESTIMATE algorithm was utilized to quantify the stromal and immune components within the endometriosis samples. The algorithm calculates characteristic ImmuneScores and StromalScores based on the gene expression matrix, providing a measure of the overall immune and stromal infiltration levels. Differential infiltration analysis of immune cell subtypes, ImmuneScores, and StromalScores between endometriosis and normal samples in the combined dataset was performed. Group comparisons were visualized using the ggplot2 package in R. Furthermore, the correlation between the expression of pyroptosis-related hub genes and the abundance of LM22 immune cells in endometriosis subtypes was assessed. Correlation analysis results were visualized using heatmaps generated with the pheatmap package in R.

### 2.11 Statistical Analysis

All statistical analyses were performed using R software (version 4.2.0). Continuous variables are presented as mean ± standard deviation. The Wilcoxon rank-sum test was used for comparisons between two groups. Categorical variables were analyzed using the chi-square test or Fisher’s exact test. Spearman correlation analysis was used to calculate correlation coefficients between different molecules. Statistical significance was defined as a p-value < 0.05 for all analyses.

## 3. Results

### 3.1 Flow Chart

Flow Chart for the Comprehensive Analysis of PRDEGs in this study is shown in Figure 1.

**Fig. 1.**
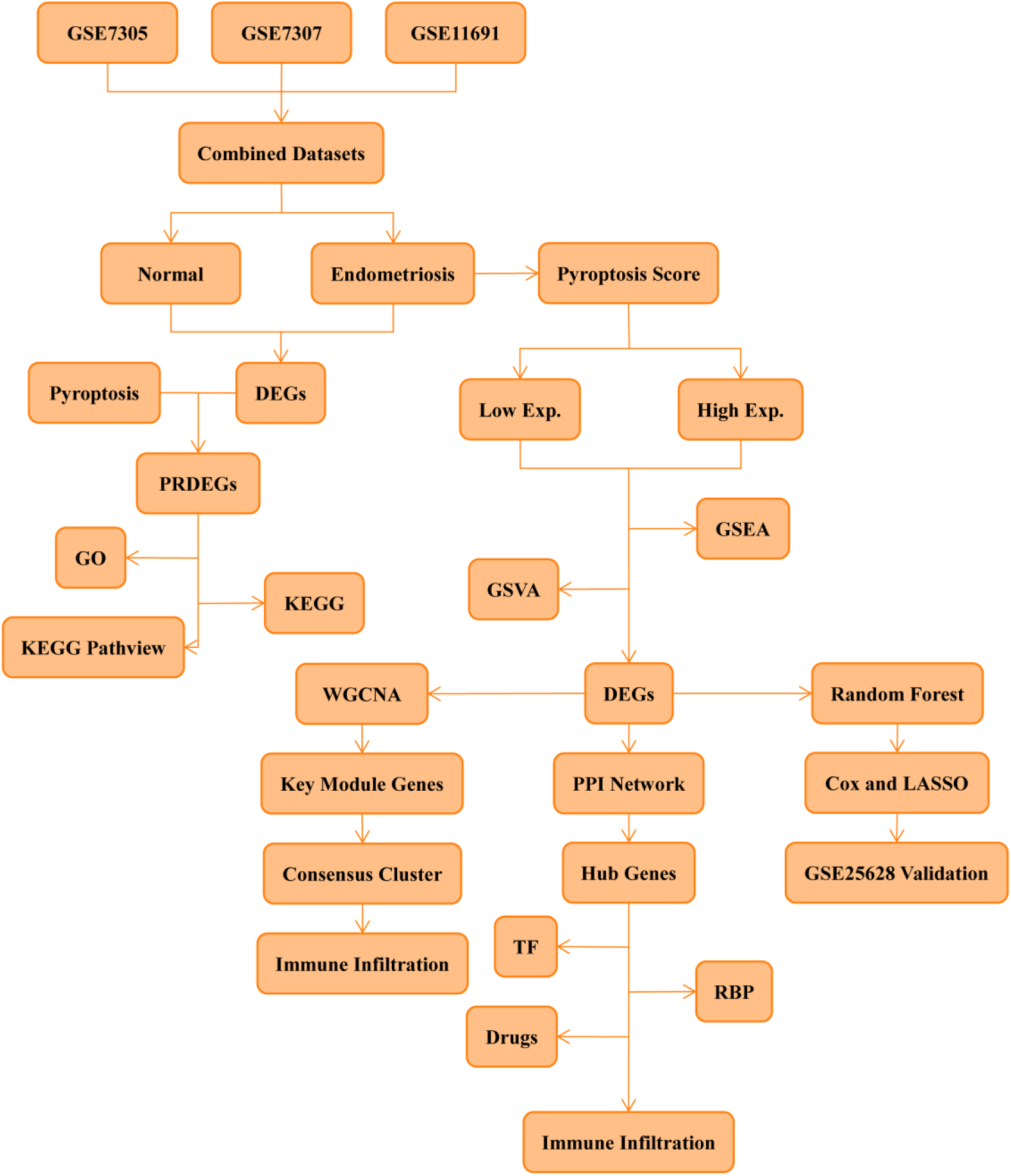
Flow Chart for the Comprehensive Analysis of PRDEGs

### 3.2 Merger of Endometriosis Datasets

To mitigate the potential influence of batch effects, the endometriosis datasets GSE7305, GSE7307, and GSE11691 were subjected to batch effect removal using the surrogate variable analysis (sva) package in R. This yielded a combined dataset for subsequent analyses. The effectiveness of batch effect removal was assessed using distribution box line plots and 3D principal component analysis (PCA) plots (Fig.2A-D). These visualizations demonstrated that the batch effect was substantially reduced following the removal process, as evidenced by the improved alignment of sample distributions and PCA clusters across the original datasets.

**Fig. 2.**
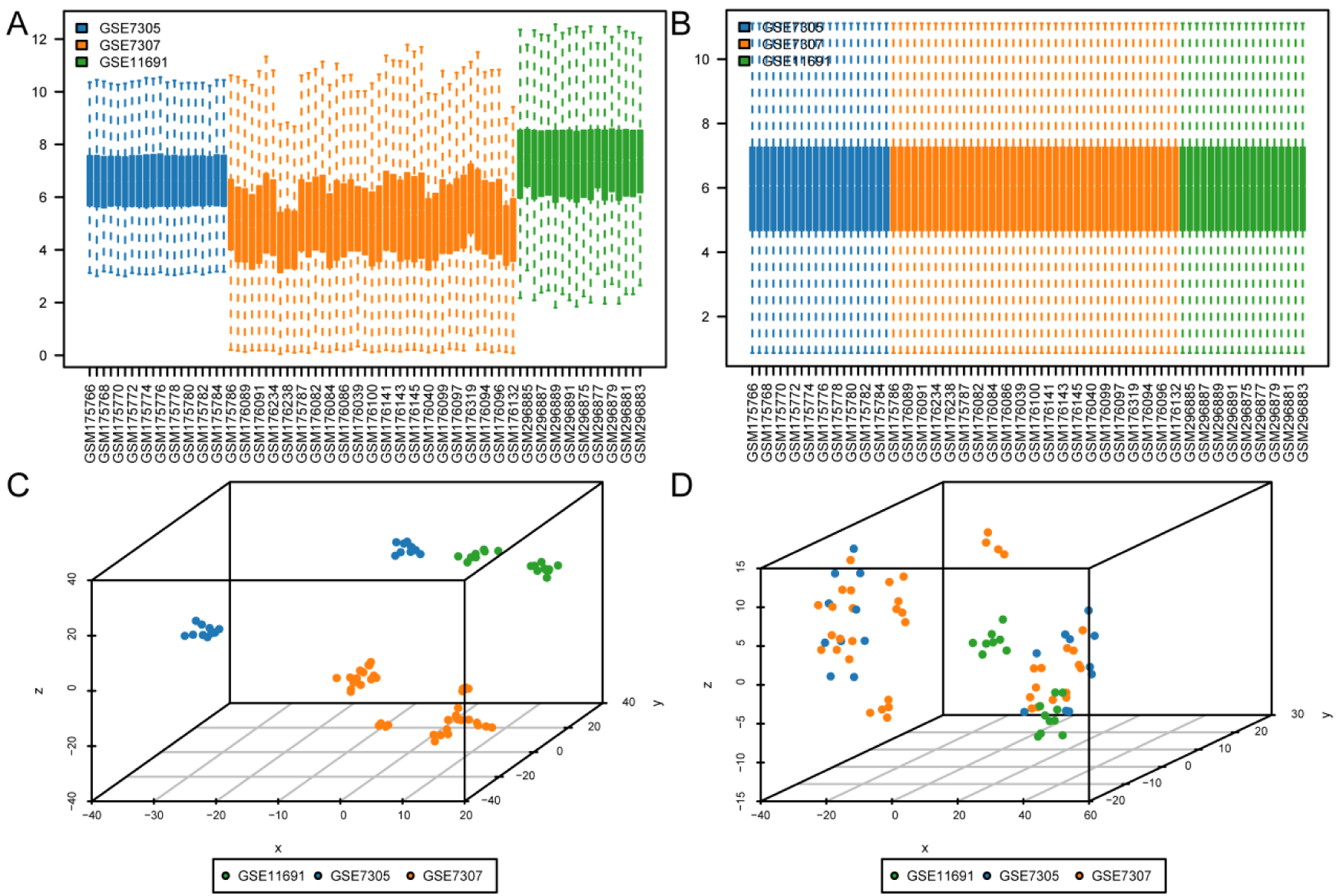
Batch Effects Removal of GSE7305, GSE7307 and GSE11691 A. Box line plots illustrating the distribution of gene expression values across datasets before batch effect removal. B. Box line plots illustrating the distribution of gene expression values in the combined dataset after batch effect removal. C. 3D PCA plot of the datasets before batch effect removal. D. 3D PCA plot of Combined Datasets after batch effect removal.

### 3.3 Identification and Characterization of PRDEGs in Endometriosis

The combined dataset was stratified into endometriosis and normal groups. Differential gene expression analysis was conducted to identify genes exhibiting significant expression changes between the two groups in the Combined Datasets. A total of 1142 DEGs were identified using a threshold of |logFC| > 1 and adjusted p-value (adj.p) < 0.05. These DEGs comprised 654 upregulated and 488 downregulated genes (Fig.3A). Subsequently, the intersection of the DEGs and the curated list of pyroptosis-related genes was determined, resulting in the identification of 26 PRDEGs (Fig.3B). These PRDEGs included VCAM1, LY96, HTRA1, BST2, FNDC4, IRAK3, CHI3L1, PPARG, SLC16A4, VDR, CD14, ICAM1, P2RX7, MKI67, APOE, PECAM1, EZH2, ADORA3, KIF23, BTK, CXCL8, CEP55, PTGS2, MELK, TREM1, and TREM2. Differential expression of the 26 PRDEGs in the Combined Dataset was visualized (Fig.3C). Finally, the chromosomal locations of these PRDEGs were mapped using the RCircos package in R, generating a chromosome localization map (Fig.3D).

**Fig. 3.**
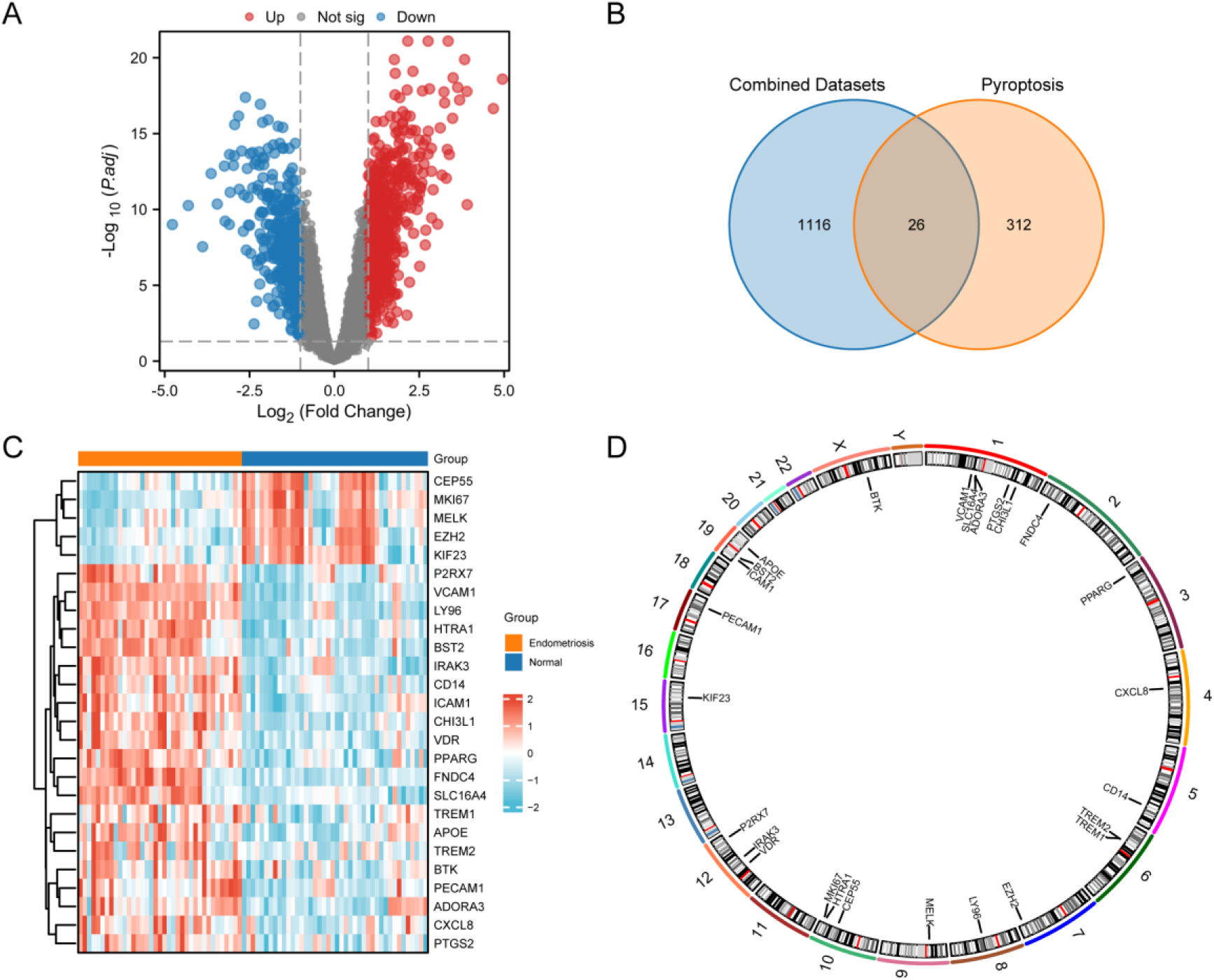
Identification and Characterization of PRDEGs in Endometriosis A. Volcano plot depicting DEGs. B. Venn diagram illustrating the overlap between DEGs and pyroptosis-related genes associated with endometriosis. C. Heat map of PRDEGs in the Combined Datasets. D. Chromosomal localization map of PRDEGs.

### 3.4 Differential Expression and Diagnostic Potential of PRDEGs in the Combined Dataset

To further validate the differential expression of the identified PRDEGs in the Combined Dataset, box line plots were generated, demonstrating significant differences in the expression levels of all 26 PRDEGs between the endometriosis and normal groups (Fig.4A). Subsequently, ROC curves were constructed for each PRDEG to assess their diagnostic potential for distinguishing between endometriosis and normal samples (Fig.4B-G). Notably, VCAM1, LY96, HTRA1, BST2, FNDC4, IRAK3, CHI3L1, PPARG, and VDR exhibited high diagnostic accuracy, with AUC values exceeding 0.9, indicating their potential as candidate diagnostic biomarkers for endometriosis.

**Fig. 4.**
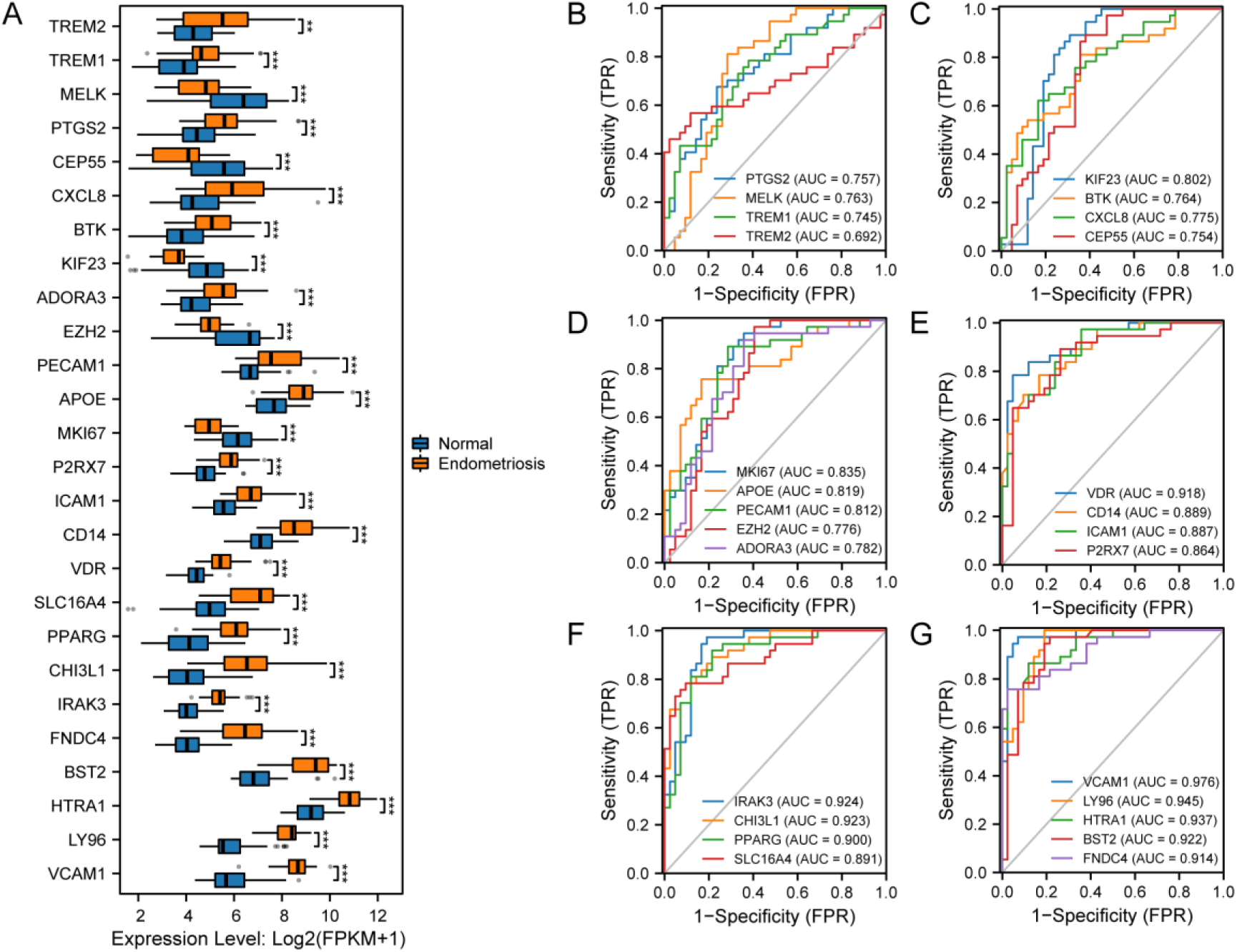
Differential Expression and Diagnostic Potential of PRDEGs in the Combined Dataset A. Box line plot illustrating the differential expression of PRDEGs in the Combined Dataset. B-G. ROC curves for 26 PRDEGs in the Combined Datasets.

### 3.5 Correlation Analysis of PRDEGs in the Combined Datasets

Utilizing the comprehensive expression matrix of 26 PRDEGs derived from the Combined Datasets, a correlation analysis was carried out (Fig. 5A). Then, the correlation analysis results between the TOP2 positively and negatively correlated genes in the correlation heat map were presented by correlation scatter plots, respectively (Fig.5B-E). The results indicated that MKI67 and MELK (r value=0.847, Fig.5B), CEP55 and MELK (r value=0.844, Fig.5C) showed a strong degree of positive correlation. MKI67 and P2RX7 (r value=-0.774, Fig.5D), MKI67 and HTRA1 (r value=-0.723, Fig.5E) showed a moderate negative correlation in Combined Datasets.

**Fig. 5.**
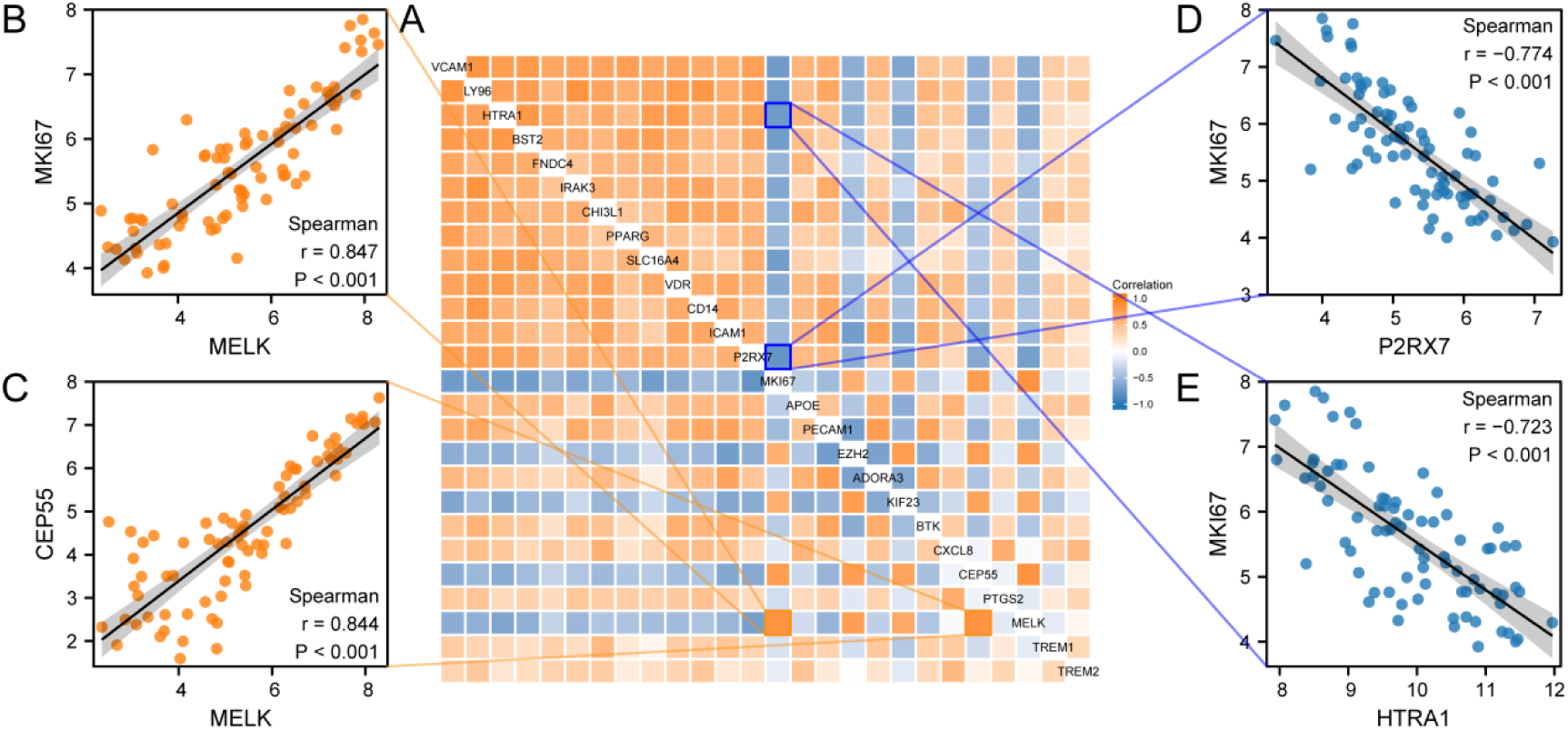
Correlation Analysis of PRDEGs A. Heatmap depicting the correlation between PRDEGs in the Combined Datasets. B-E. Scatter plots illustrating the correlations between: MKI67 and MELK (B), CEP55 and MELK (C), MKI67 and P2RX7 (D), and MKI67 and HTRA1 (E).

### 3.6 GO and KEGG Enrichment Analysis of PRDEGs

To elucidate the biological functions and pathways associated with the 26 PRDEGs, GO and KEGG enrichment analyses were performed. The results of these analyses are visualized in Figure 6A and detailed in Table S2. Network diagrams were constructed to illustrate the relationships between enriched GO terms within the BP, CC, and MF categories, as well as enriched KEGG pathways (Fig. 6B-E). Furthermore, a combined logFC analysis of GO and KEGG enrichment was conducted for the 26 PRDEGs (Fig.6F). Notably, the visualized histogram of the GO and KEGG enrichment analyses suggests that the NF-kappa B signaling pathway may be the most significantly positively regulated pathway, while the Flemming body appears to be the most prominently negatively regulated pathway.

**Fig. 6.**
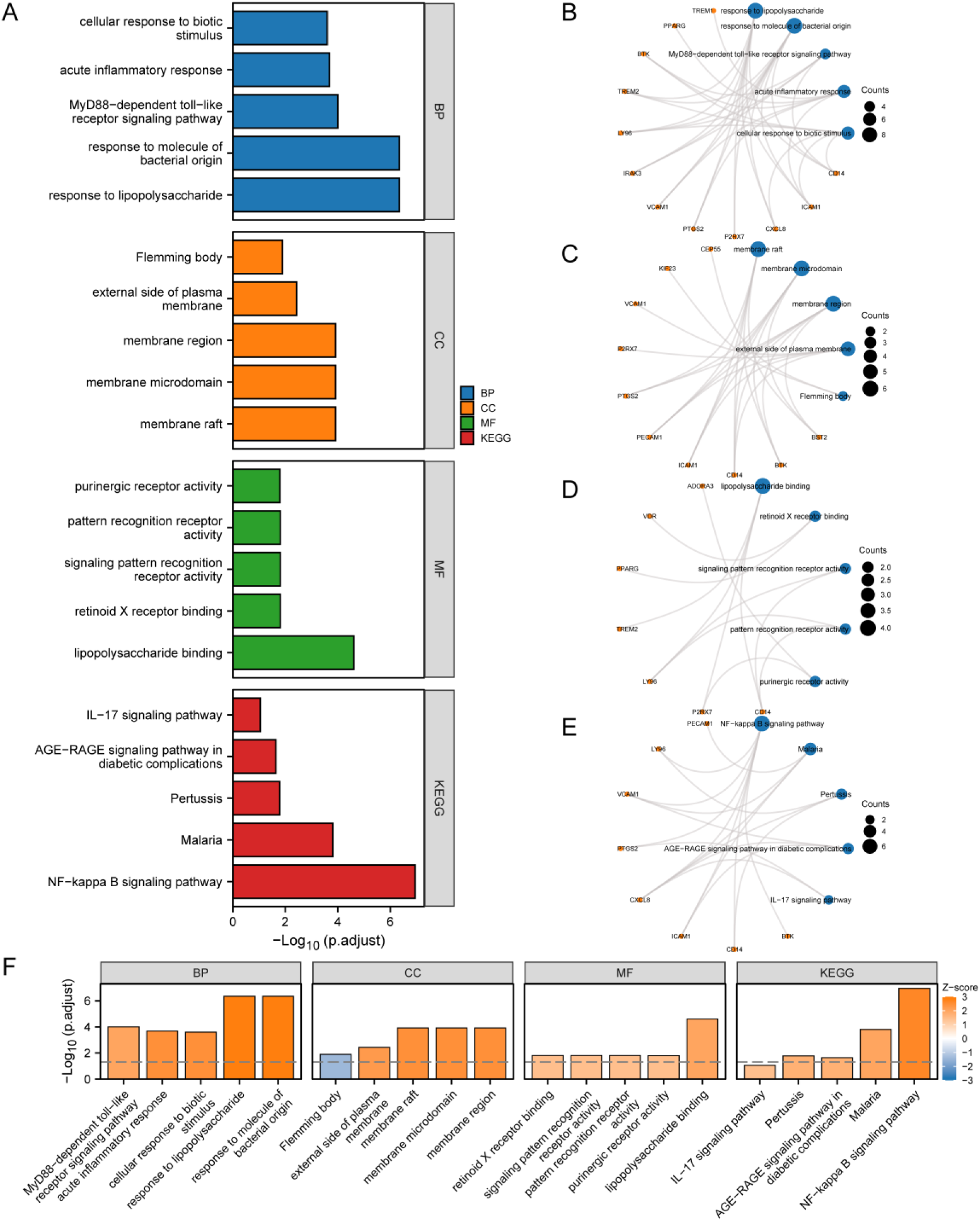
GO and KEGG Enrichment Analysis of PRDEGs A. Histogram of GO and KEGG enrichment analysis of PRDEGs. B-E. Network diagrams of GO and KEGG enrichment analysis. Blue nodes represent entries, orange nodes represent molecules, and connecting lines represent the relationship between terms/pathways and genes. F. Combined logFC histogram.

To further investigate the involvement of specific pathways identified through KEGG enrichment analysis, pathway maps were generated for the IL-17 signaling pathway (Fig.7A), Pertussis (Fig.7B), Malaria (Fig.7C), and the NF-kappa B signaling pathway (Fig.7D).

**Fig. 7.**
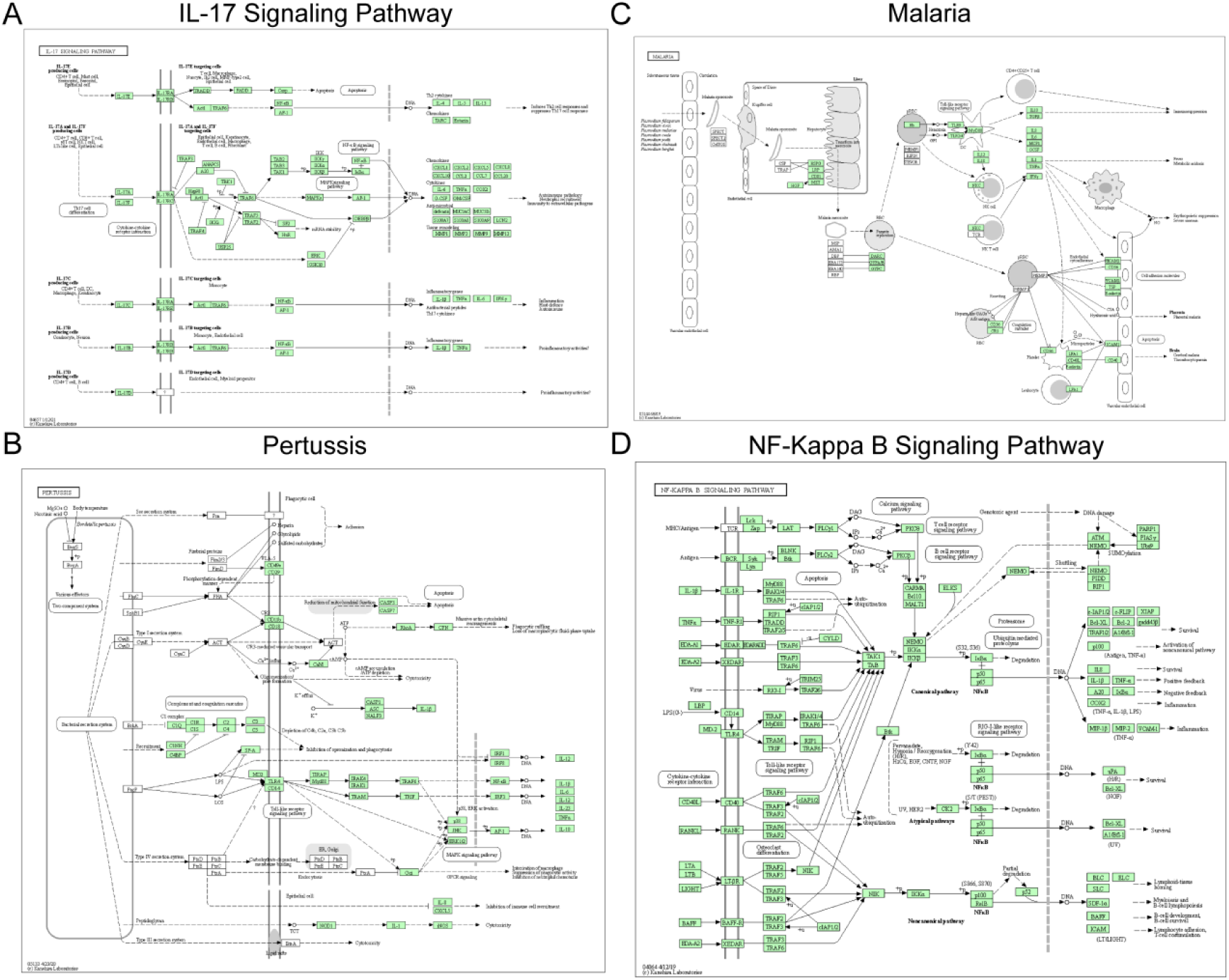
KEGG Pathway Visualization A-D. Pathway map for KEGG enrichment analysis of PRDEGs: IL-17 signaling pathway (A), Pertussis (B), Malaria (C) and NF-kappa B signaling pathway (D).

### 3.7 Pyroptosis Score Analysis and Differential Gene Expression between High and Low Pyroptosis Expression Groups in Endometriosis

A pyroptosis score was calculated for each endometriosis patient in the combined dataset based on the expression levels of the 26 PRDEGs. Patients were then stratified into high and low pyroptosis expression groups based on the median pyroptosis score. A violin plot (Fig.8A) revealed a statistically meaningful difference in pyroptosis scores between the two groups (p < 0.001). Furthermore, the ROC curve (Fig.8B) demonstrated that the pyroptosis score exhibited excellent discriminatory power for distinguishing between the two groups, with an AUC of 1. Additionally, differential gene expression analysis was performed between the two groups. A total of 238 DEGs were recognized using a threshold of |logFC| > 0 and adjusted p-value (adj.p) < 0.05, comprising 203 upregulated and 35 downregulated genes (Fig.8C).

**Fig. 8.**
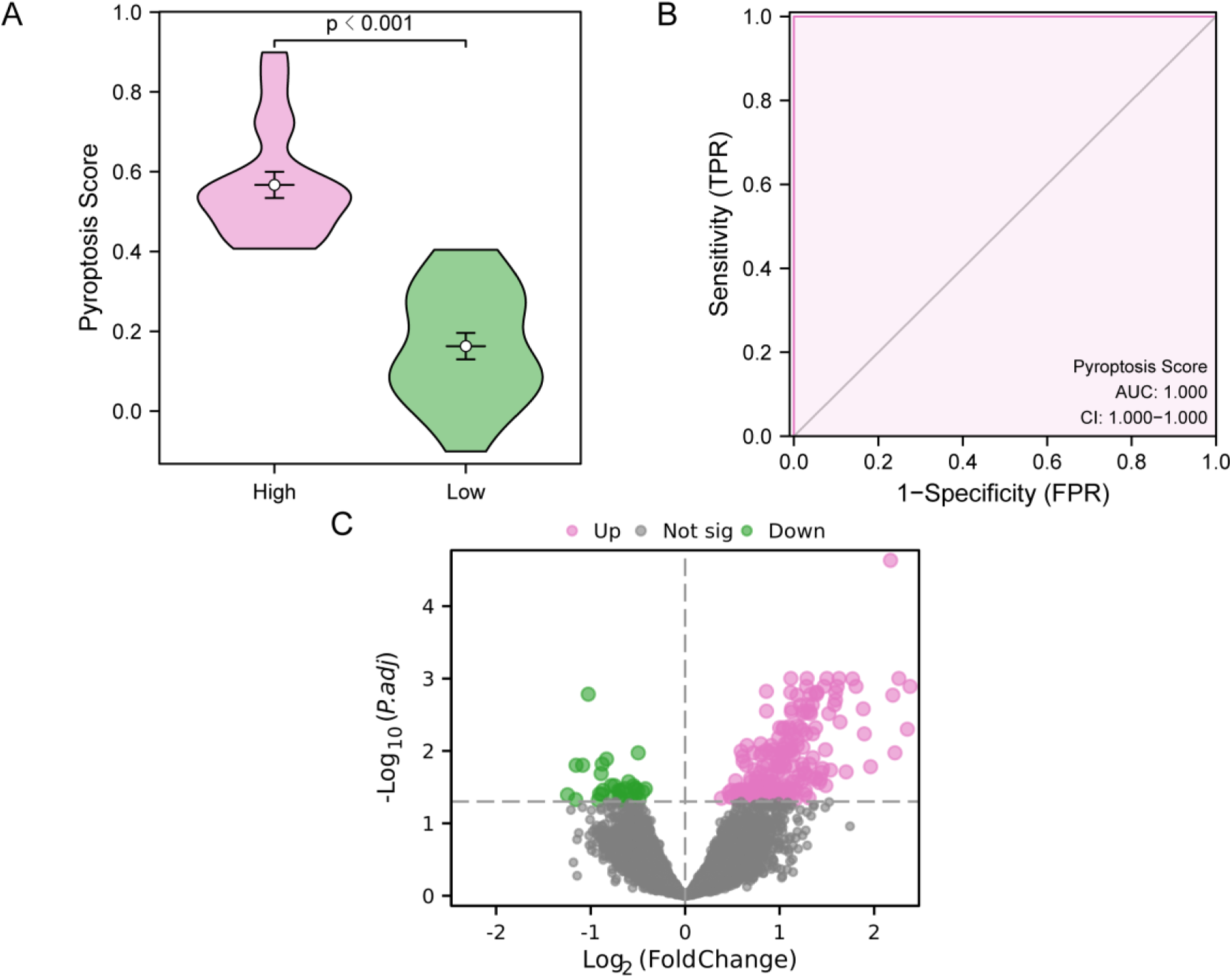
Differential Gene Expression Analysis Based on Pyroptosis Score A. Violin plot illustrating the distribution of pyroptosis scores in the subgroups. B. ROC curve for the pyroptosis score, demonstrating its ability to discriminate between the two subgroups. C. Volcano plot depicting DEGs.

### 3.8 Development and Evaluation of a Diagnostic Model for Endometriosis based on DEGs

To evaluate: the diagnostic potential of the 238 DEGs identified between the high and low pyroptosis expression groups in endometriosis, the Random Forest algorithm was employed (Fig.9A). This analysis identified 123 DEGs with an IncNodePurity score greater than 0.2, indicating their importance in differentiating between the two groups. The top 20 DEGs ranked by IncNodePurity are shown in Fig. 9B. Thereafter, a multifactorial Cox regression model was formed using these DEGs. The resulting model included 15 DEGs: KIF13B, SRGAP3, BAG6, MYO5A, GPKOW, TRAP1, HEATR2, SIGLEC1, RNF34, MGAT4A, EDNRA, CTSZ, NPM3, AK055981, and TLR8. Furthermore, a diagnostic model for endometriosis was developed using LASSO regression (Fig.9C). The trajectory of LASSO variable selection is visualized in Fig.9D. The final LASSO model included 5 DEGs: KIF13B, BAG6, MYO5A, HEATR2, and AK055981 (Fig.9E). The prognostic performance of this diagnostic model was evaluated using nomogram analysis (Fig.9F).

**Fig. 9.**
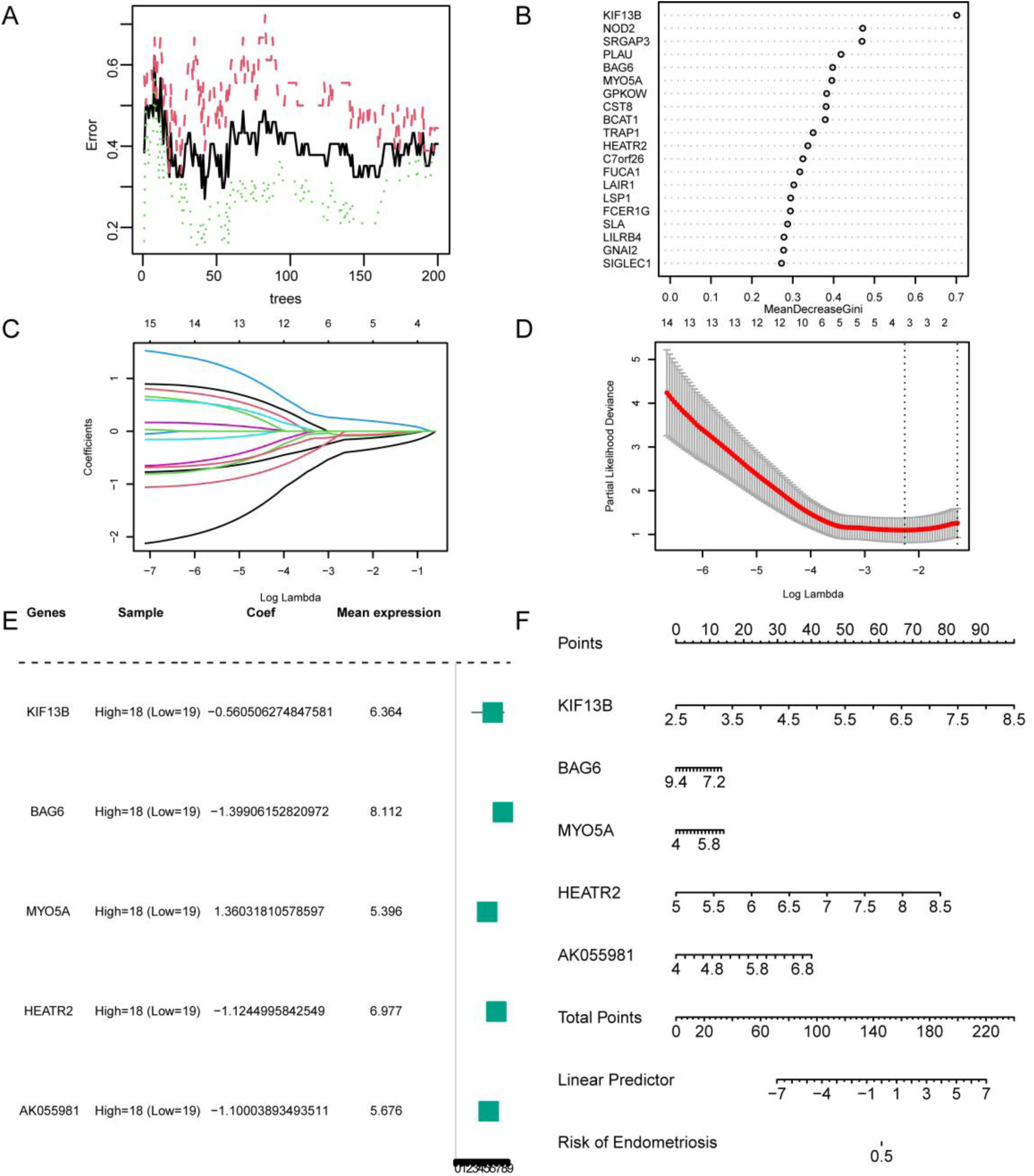
Random Forest, Cox and LASSO Regression Analysis of the diagnostic model A. Plot of the training error for the Random Forest algorithm during model development. B. Top 20 DEGs identified by the Random Forest model. C. Diagnostic model plot based on 15 DEGs identified through multifactorial Cox regression analysis. D.Trajectory plots illustrating the prognostic value of the 15 DEGs identified through multifactorial Cox regression analysis. E. Forest plot depicting the hazard ratios and confidence intervals for the 5 DEGs selected by LASSO regression analysis.

### 3.9 Validation of the diagnostic model of Endometriosis

Calibration analysis was performed to assess the accuracy and discriminative ability of the developed diagnostic model for endometriosis (Fig.10A). The clinical utility of the model was further evaluated using DCA plots (Fig.10B). These analyses demonstrated that the diagnostic model exhibited good calibration and provided a net benefit across a range of threshold probabilities. To validate the generalizability of the model, the LASSO scores for pyroptosis-related LASSO genes were calculated in the independent validation dataset GSE25628 based on the LASSO coefficients derived from the training dataset. Calibration curves and DCA plots were then generated for the validation dataset using the 5 DEGs included in the diagnostic model (Fig.10C-D). The results confirmed that the diagnostic model maintained its accuracy and clinical utility in the independent validation cohort, suggesting its robustness and potential for broader application.

**Fig. 10.**
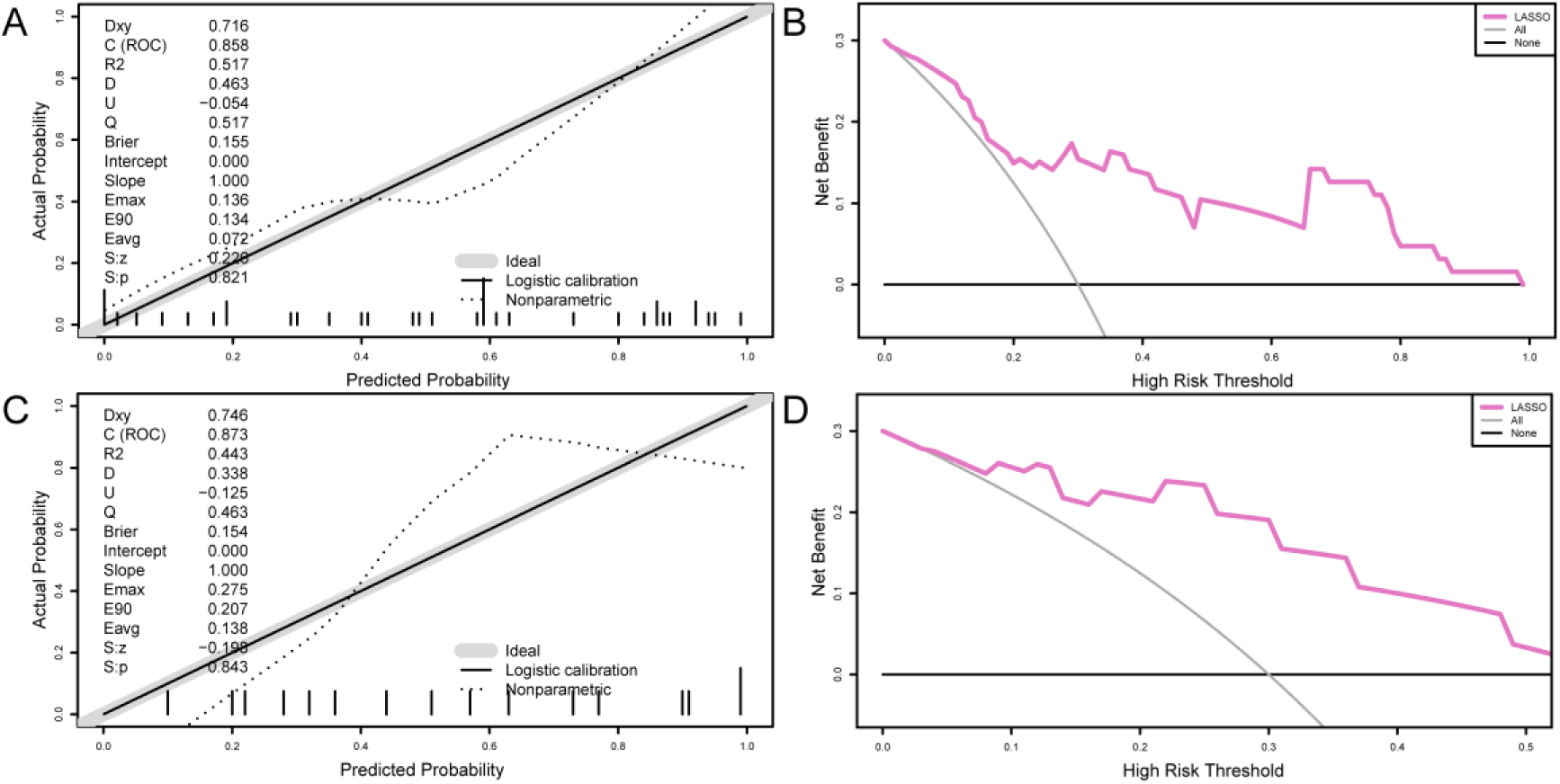
Module Validation and Datasets Validation of GSE25628 A-B: Calibration curve(A), DCA plots(B) for the diagnostic model based on 5 DEGs in the training dataset. C-D. Calibration curve(C), DCA plots(D) for the diagnostic model in the independent validation dataset GSE25628.

### 3.10 GSEA of Endometriosis-Related Genes in the Combined Dataset

GSEA was performed to investigate the biological functions and pathways associated with gene expression changes in the endometriosis group of the combined dataset,. This analysis examined the relationship between the expression levels of all genes and curated gene sets representing BP, CC, and MF. The results of the GSEA are visualized in Fig.11A and detailed in Table S3. The GSEA revealed that the expression of genes in the endometriosis group markedly impacted several biologically relevant functions and signaling pathways. Remarkably, enriched pathways included those related to IL1 and megakaryocytes in obesity (Fig.11B), the IL17 pathway (Fig.11C), the IL12 pathway (Fig.11D), and the IL3 signaling pathway (Fig.11E), etc.

**Fig. 11.**
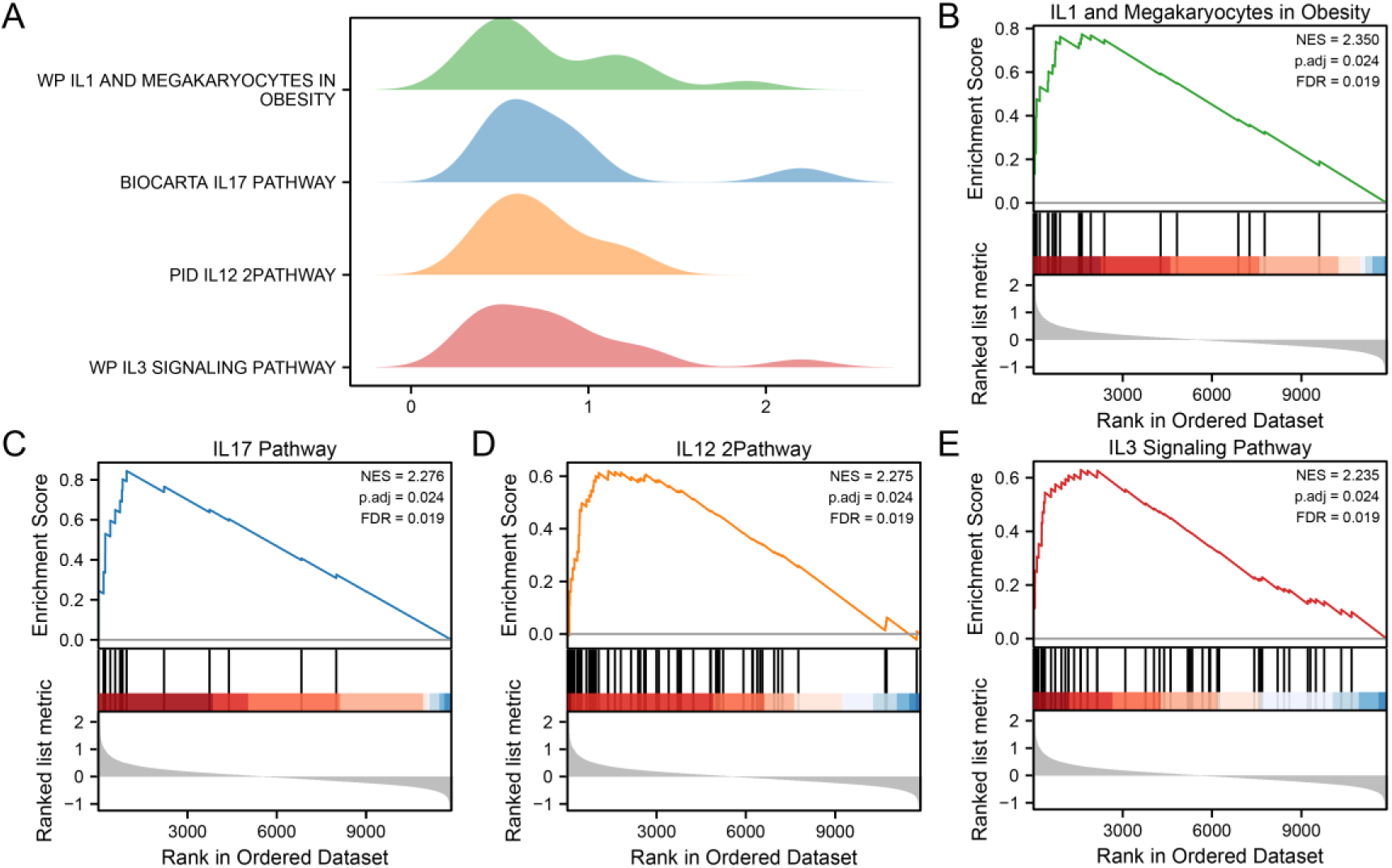
GSEA of Endometriosis-Related Genes in the Combined Dataset A. Enrichment plots for the top 4 biological functions enriched in the endometriosis group of the combined dataset. B-E. GSEA enrichment plots demonstrating the significant impact of endometriosis on the following pathways: IL1 and Megakaryocytes in Obesity (B), IL17 Pathway (C), IL12 2Pathway (D) and IL3 Signaling Pathway (E).

### 3.11 GSVA of Hallmark Pathways in Endometriosis Subgroups

To investigate the differential enrichment of hallmark pathways between the high and low pyroptosis expression groups in endometriosis, GSVA was performed using the expression data of all genes in the combined dataset. The results of the GSVA are detailed in Table S4. Hallmark pathways exhibiting a meaningful difference in enrichment between the two groups (p-value < 0.05) were identified and visualized using a grouped comparison box plot (Fig.12A). The GSVA revealed that 15 hallmark pathways were appreciably enriched in either the high or low pyroptosis expression group (p < 0.05). Finally, the distinctive enrichment of these 15 hallmark pathways between the high and low pyroptosis expression groups was visualized using a heatmap (Fig.12B).

**Fig. 12.**
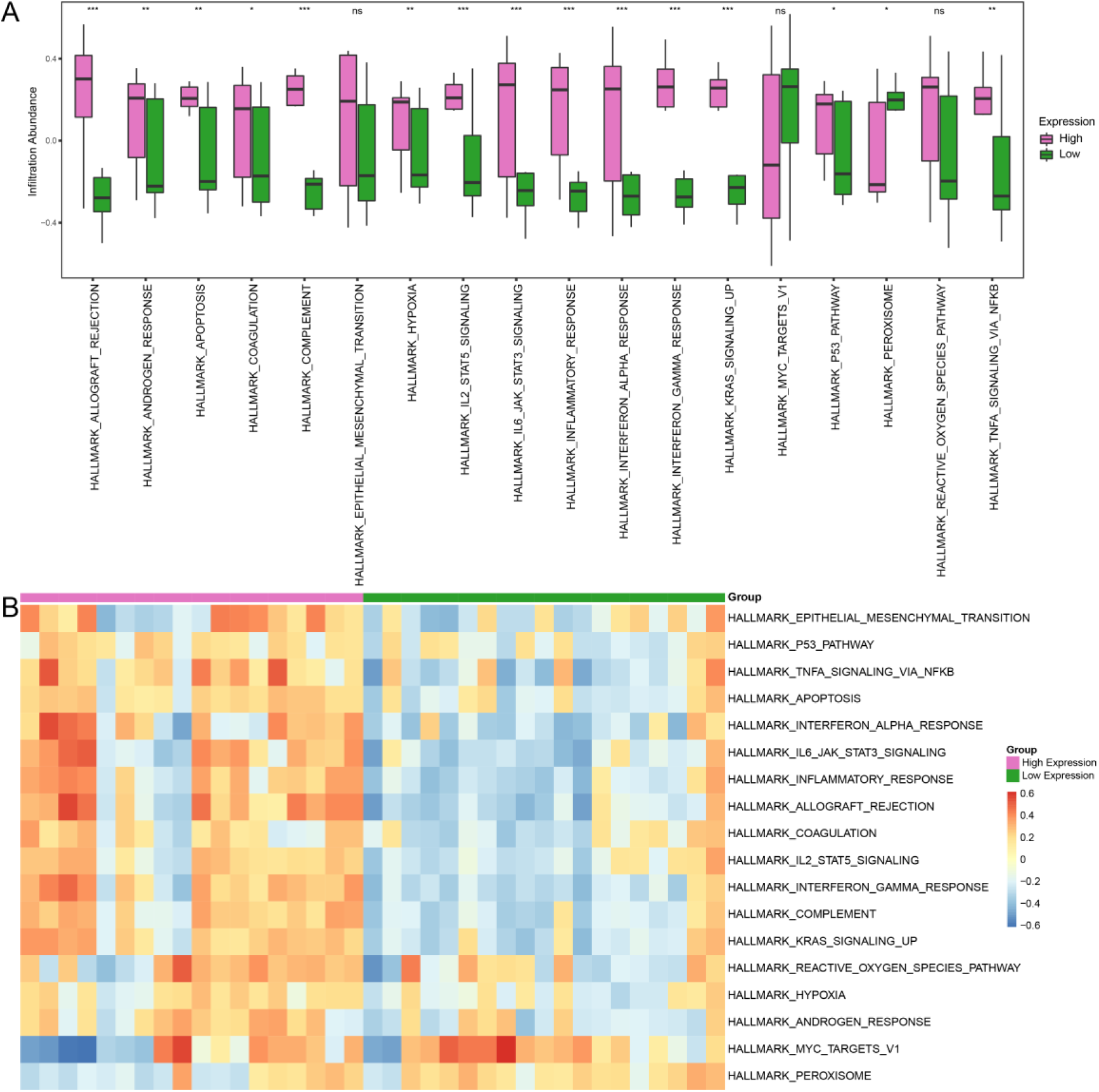
GSVA of Hallmark Pathways in Endometriosis Subgroups A. Box plots illustrating the differential enrichment of hallmark pathways between the high and low pyroptosis expression groups. B. Heatmap depicting the enrichment scores of hallmark pathways in the two groups.

### 3.12 WGCNA and Identification of KMGs in Endometriosis Subgroups

To identify co-expression modules within the endometriosis group of the combined dataset, weighted gene WGCNA was performed using the genes exhibiting the highest 25% variance. Initially, the scale-free fit index was calculated under individual soft thresholding powers (Fig.13A). An optimal soft thresholding power of 3 was selected, corresponding to a scale-free fit index of 0.90. Using the optimal soft thresholding power, a co-expression network was constructed, and DEGs were clustered using a hierarchical clustering tree (Fig.13B). With a module merging threshold of 0.2, the genes in the top 25% variance range were clustered into six distinct modules: MEblue, MEyellow, MEturquoise, MEgreen, MEbrown, and MEred. The relationship between these genes and their assigned modules is illustrated in Fig.13C. The correlation between the six module eigengenes and the high and low pyroptosis expression subgroups of endometriosis was then analyzed (Fig.13D). The MEturquoise (|r value| = 0.48) and MEgreen (|r value| = 0.45) modules exhibited the strongest correlations with the pyroptosis expression subgroups and were selected for further analysis. Finally, the DEGs identified between the high and low pyroptosis expression subgroups were intersected with the genes within the MEturquoise (Fig.13E) and MEgreen (Fig.13F) modules, respectively, using Venn diagrams. This analysis yielded a total of 21 key module genes (KMGs): NOD2, IL32, AGRP, ASGR2, BCAT1, COL11A1, SLC12A8, TLR8, MGAT4A, NABP1, SERPINE1, SAMHD1, GDF2, KIF13B, NEUROD4, CST8, NLGN4Y, UBASH3A, TNF, HSD11B2, and LOC100131510.

**Fig. 13.**
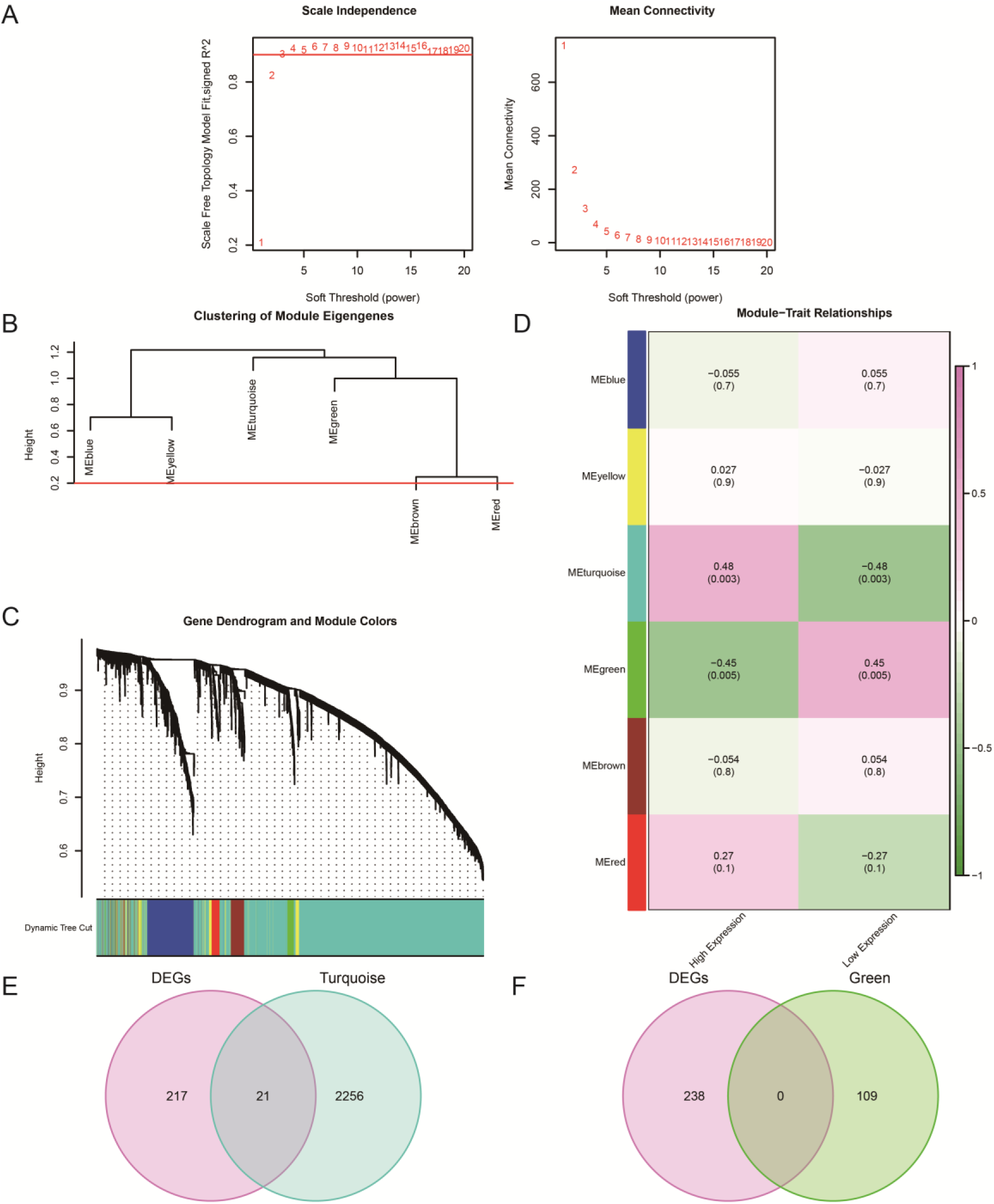
WGCNA and Identification of KMGs A. Scale-free network analysis for determining the optimal soft thresholding power in WGCNA. The left panel shows the scale-free fit index, and the right panel depicts the mean network connectivity. B. Dendrogram illustrating the hierarchical clustering of gene co-expression modules for genes exhibiting the top 25% variance. C. Module assignment for genes within the top 25% variance range. The upper panel shows the hierarchical clustering dendrogram, and the lower panel displays the corresponding module colors assigned to each gene. D. Correlation analysis between module eigengenes and the high and low pyroptosis expression groups. E-F. Venn diagrams depicting the overlap between DEGs and genes within the:(E) MEturquoise module, (F) MEgreen module.

### 3.13 PPI Network Analysis and Identification of Hub Genes

To investigate the interactions among the 21 KMGs, a PPI network was constructed using the STRING database. The resulting network comprising 13 interacting KMGs was visualized (Fig.14A). The interacting KMGs included NOD2, IL32, AGRP, ASGR2, BCAT1, COL11A1, TLR8, SERPINE1, SAMHD1, GDF2, NLGN4Y, UBASH3A, TNF, HSD11B2, and LOC100131510.

**Fig. 14.**
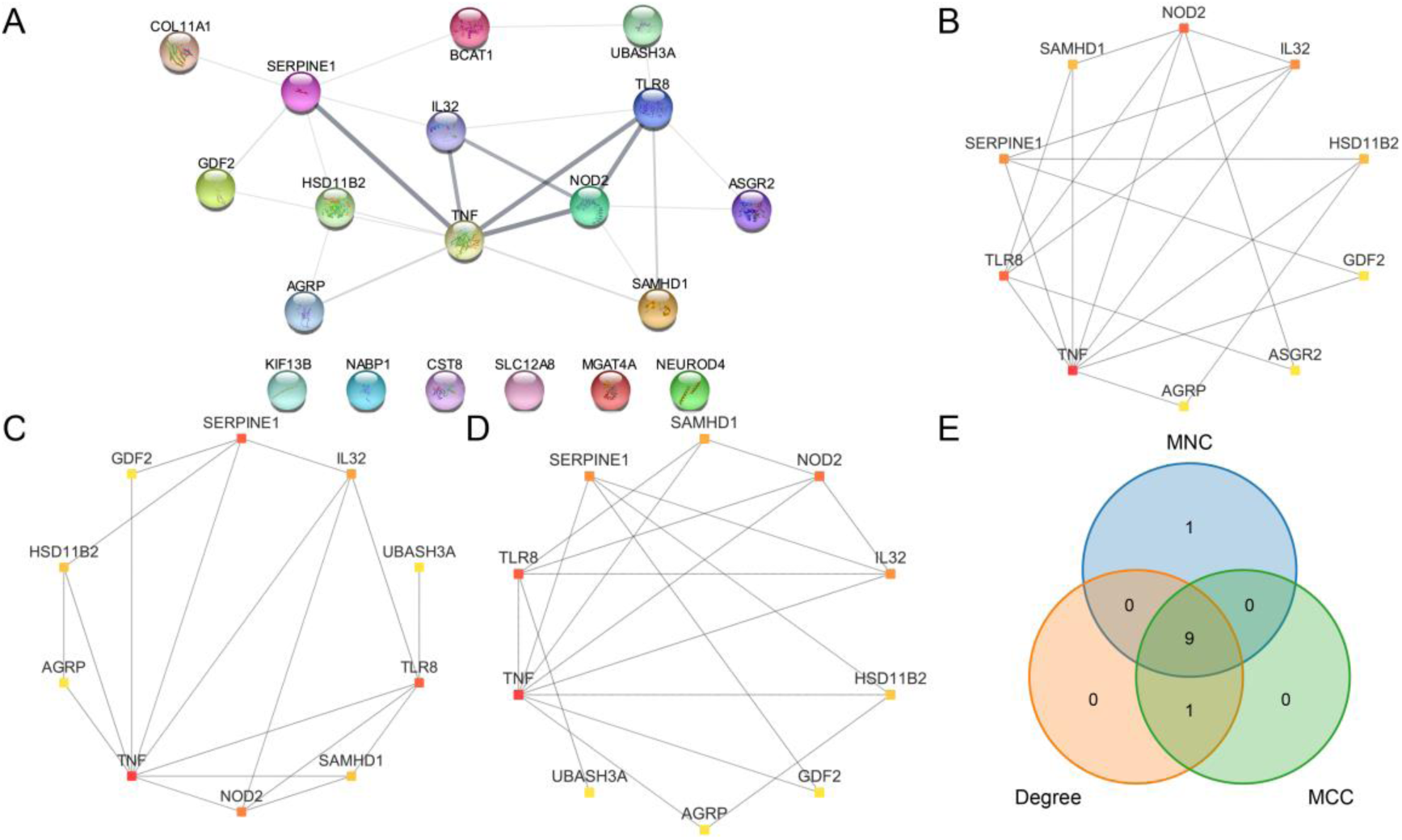
Network Analysis and Identification of Hub Genes A. PPI network of KMGs. B. Top 10 KMGs ranked by MNC algorithm. C. Top 10 KMGs ranked by Degree algorithm. D. Top 10 KMGs ranked by MCC algorithm. E. Venn diagram illustrating the overlap among the top 10 KMGs identified by the MNC, Degree, and MCC algorithms. Node colors represent the ranking of each gene, with red indicating the highest rank and yellow indicating the lowest rank.

Subsequently, three algorithms in Cytoscape were employed to identify hub genes within the PPI network: Maximum Neighborhood Component (MNC) (Fig.14B), Degree (Fig.14C), and Maximal Clique Centrality (MCC) (Fig.14D). The top 10 genes ranked by each algorithm were then intersected and visualized (Fig.14E). It identified 9 pyroptosis-related hub genes: TNF, NOD2, TLR8, IL32, SERPINE1, SAMHD1, HSD11B2, GDF2, and AGRP.

### 3.14 Construction of Regulatory Networks Involving Pyroptosis-Related Hub Genes

To explore the potential regulatory roles of miRNAs and circRNAs in pyroptosis, miRNA-hub gene interactions and circRNA-miRNA interactions were retrieved from the StarBase database. This analysis yielded a network comprising 6 pyroptosis-related hub genes, 28 miRNAs, and 44 circRNAs. The resulting ceRNA network was constructed (Fig.15A). Detailed information regarding these interactions is provided in Table S5.

**Fig. 15.**
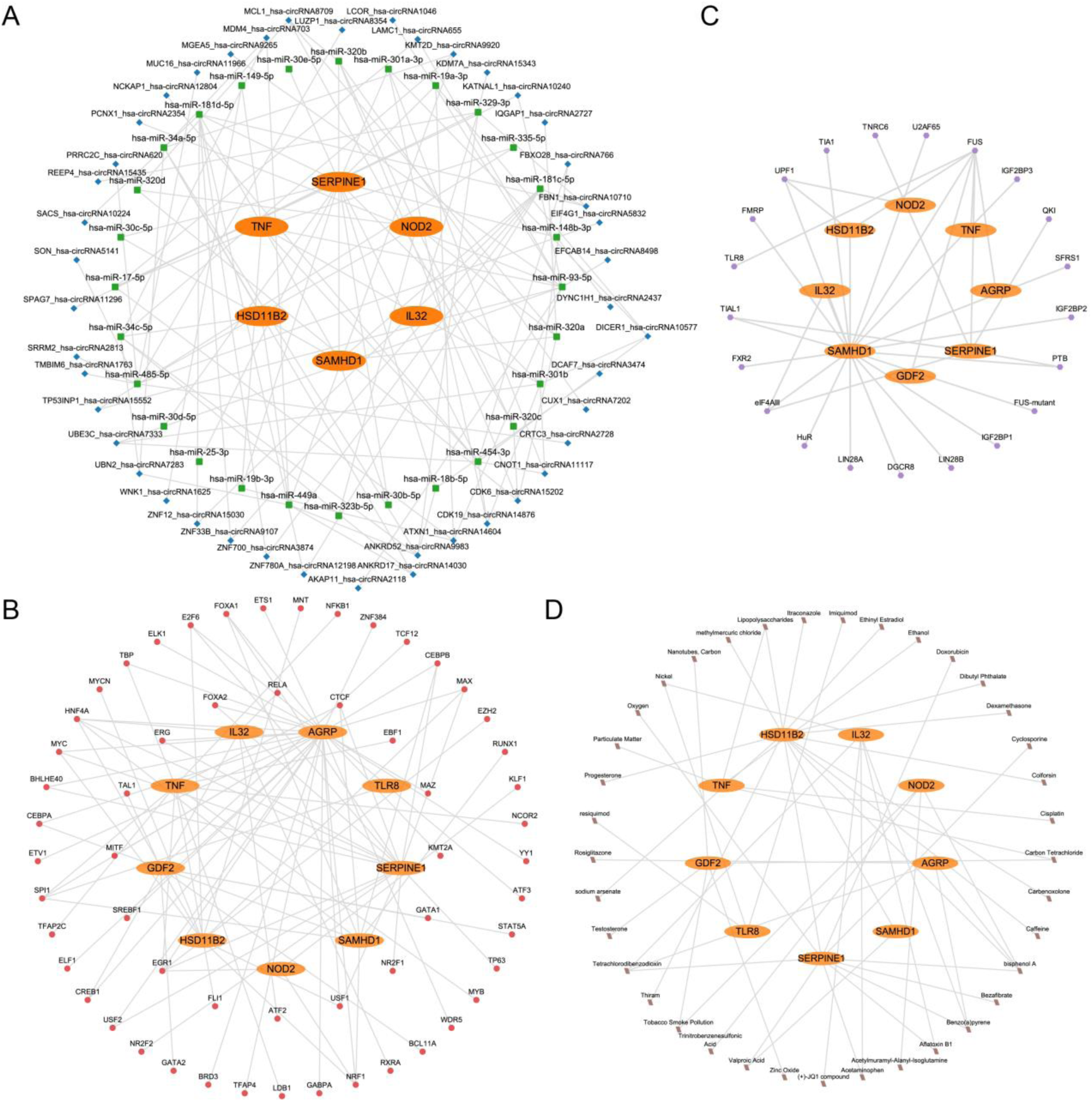
ceRNA and Regulatory Network of Hub Genes. A-D. ceRNA Network(A), TF-mRNA Regulatory Network(B), mRNA-RBP Regulatory Network(C), mRNA-Drugs Regulatory Network(D) of pyroptosis-related hub genes. orange oval for mRNA, green square for miRNA, blue diamond for circRNA, purple hexagon for TF, red circle for RBP, brown parallelogram for Drugs.

To investigate the transcriptional regulation of pyroptosis-related hub genes, TF-hub gene interactions were characterized using the ChIPBase database. A TF-mRNA regulatory network was then constructed, encompassing 9 pyroptosis-related hub genes and 57 TFs (Fig.15B). Detailed information regarding these interactions is provided in Table S6.

To explore the post-transcriptional regulation of pyroptosis-related hub genes, potential RNA-binding protein (RBP)-hub gene interactions were predicted. An mRNA-RBP regulatory network was subsequently constructed, comprising 9 pyroptosis-related hub genes and 21 RBPs (Fig.15C). Detailed information regarding these interactions is provided in Table S7.

To investigate potential pharmacological agents or molecular compounds targeting pyroptosis-related hub genes, the Comparative Toxicogenomics Database (CTD) was queried. A drug-mRNA regulatory network was constructed, encompassing 9 pyroptosis-related hub genes and 37 drugs or molecular compounds (Fig.15D). Detailed information regarding these interactions is provided in Table S8.

### 3.15 Immune Infiltration Landscape in Endometriosis

To characterize the immune microenvironment in endometriosis, the abundance of immune cell infiltration was assessed in both endometriosis and normal endometrial samples using the CIBERSORT, ESTIMATE, and ssGSEA algorithms. The CIBERSORT analysis identified significant differences in the composition of 11 immune cell types across endometriosis and normal subgroups (Fig. 16A, p < 0.05). Notably, macrophages M2, activated mast cells, activated NK cells, and plasma cells exhibited particularly pronounced differences (p < 0.001). The ESTIMATE algorithm, which quantifies the overall immune and stromal content, demonstrated significant differences in the Estimate Score, ImmuneScore, and StromalScore between endometriosis and normal samples (Fig.16B, p < 0.001). These findings suggest a distinct stromal and immune landscape in endometriosis compared to normal endometrium. The ssGSEA analysis, which assesses the enrichment of immune cell-specific gene sets, identified 21 immune cell types with significantly different enrichment scores between endometriosis and normal subgroups (Figure 16C, p < 0.05).

**Fig. 16.**
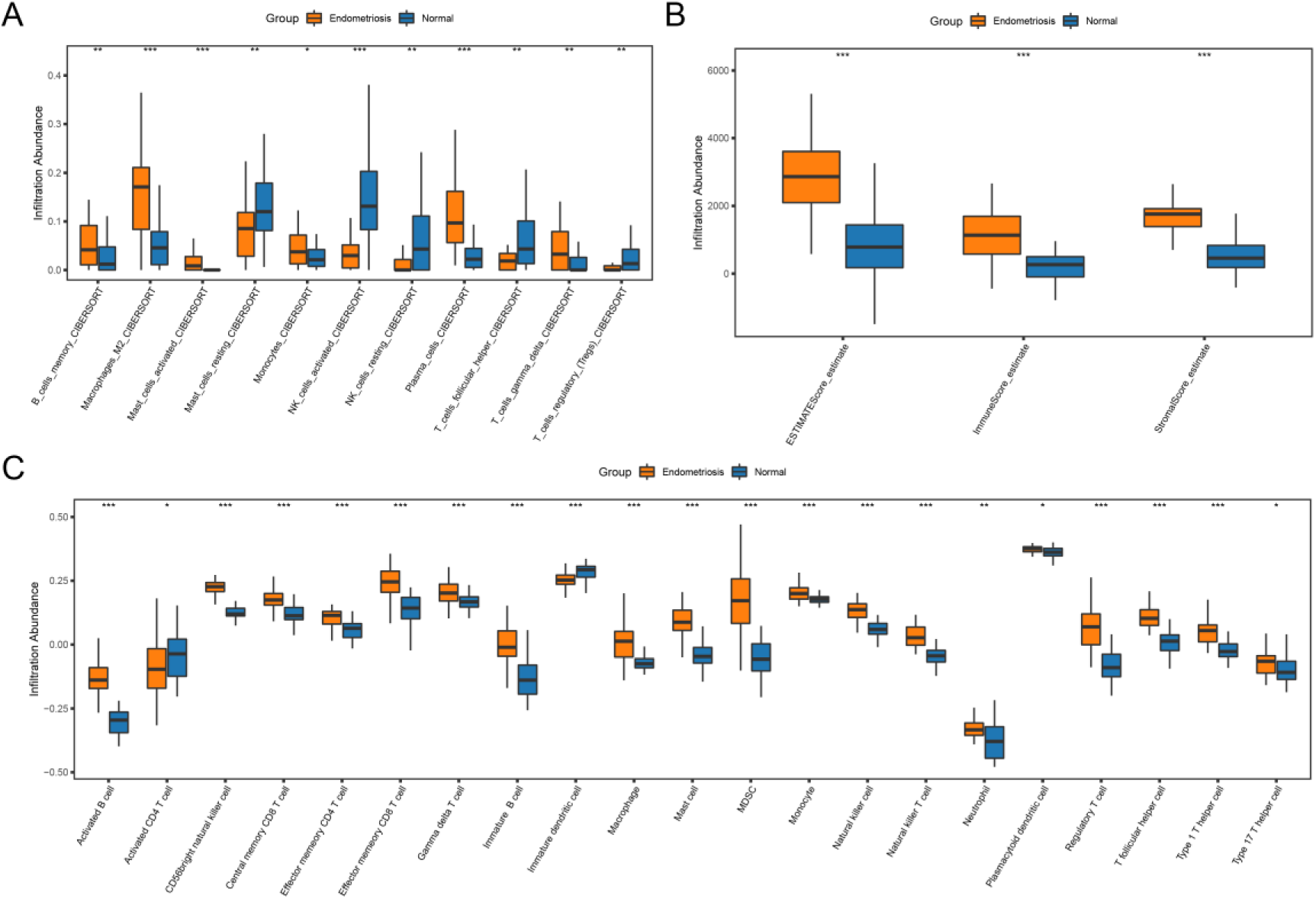
Immune Infiltration Landscape in Endometriosis A. Box plots comparing the abundance of immune cell subtypes identified by CIBERSORT between endometriosis and normal subgroups. B. Box plots comparing the ESTIMATE scores (Estimate Score, ImmuneScore, and StromalScore) between endometriosis and normal subgroups. c. Box plots comparing the enrichment scores of immune cell types identified by ssGSEA between endometriosis and normal subgroups.

### 2.16 Identification of Endometriosis Subtypes Based on KMGs Expression

To explore the heterogeneity of endometriosis based on KMG expression, the ConsensusClusterPlus package in R was employed to classify endometriosis samples into distinct subtypes using the expression profiles of the 21 KMGs. This analysis identified three distinct endometriosis subtypes, designated as Cluster 1 (n = 12 samples), Cluster 2 (n = 9 samples), and Cluster 3 (n = 16 samples) (Fig.17A-B). Principal component analysis (PCA) plots confirmed that these three subtypes exhibited distinct gene expression patterns (Fig.17C). A heatmap was generated to visualize the differential expression of the 21 KMGs across the three endometriosis subtypes (Fig.17D). Furthermore, Subtype-specific KMG expression levels showed significant differences, as illustrated by a violin plot (Fig.17E, p < 0.05).

**Fig. 17.**
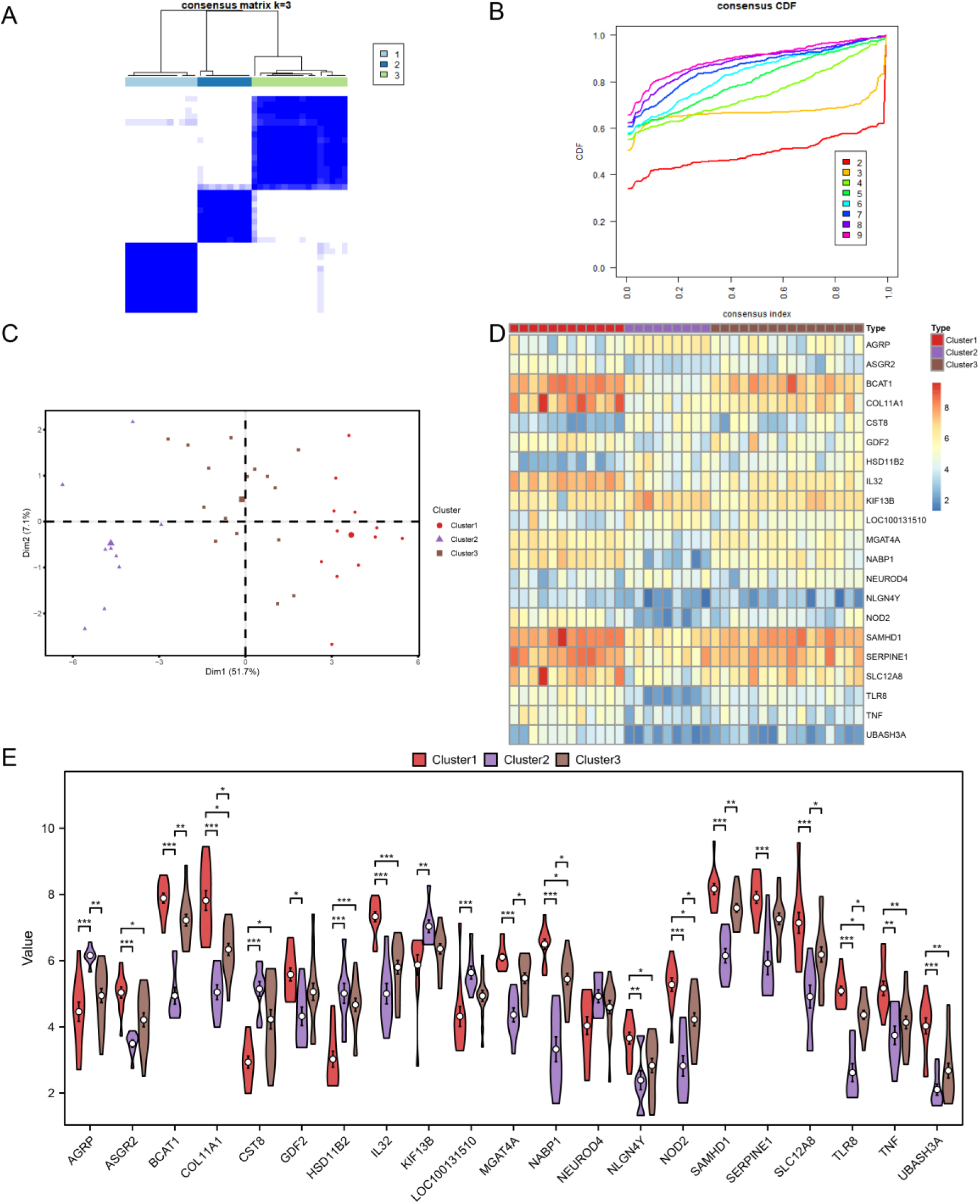
Identification of Endometriosis Subtypes Based on KMG Expression A. Consensus clustering matrix for endometriosis samples, demonstrating the stability of the identified subtypes. B. Consistent Cumulative Distribution Function (CDF) plot for the consensus clustering analysis, indicating the optimal number of clusters. C. PCA plot visualizing the separation of the three identified endometriosis subtypes. D. Heatmap depicting the expression levels of KMGs across the three endometriosis subtypes. E. Violin plots of KMG expression levels within each of the three endometriosis subtypes.

### 2.17 Correlation Analysis of Immune Cell Infiltration and KMG Expression in Endometriosis Subtypes

The relationship between immune cell infiltration and KMG expression in endometriosis subtypes was investigated by correlating the expression of 22 immune cell types with the three subtypes using the endometriosis group’s expression data. A histogram was generated to visualize the proportion of each immune cell type within each endometriosis subtype (Fig.18A). Subsequently, the correlation between immune cell infiltration abundance and endometriosis subtypes was analyzed (Fig. 18B-D). The correlation heatmaps revealed distinct patterns of immune cell infiltration across the subtypes. In Cluster 1 (Fig.18B), T follicular helper cells and gamma delta T cells exhibited the strongest positive correlation, while activated mast cells and resting mast cells showed a strong negative correlation. In Cluster 2 (Fig.18C), activated mast cells and plasma cells demonstrated the strongest positive correlation, whereas resting dendritic cells and memory B cells exhibited the strongest negative correlation. In Cluster 3 (Fig.18D), activated mast cells and T follicular helper cells showed the strongest positive correlation, while M1 macrophages and activated NK cells displayed the most pronounced negative correlation.

**Fig. 18.**
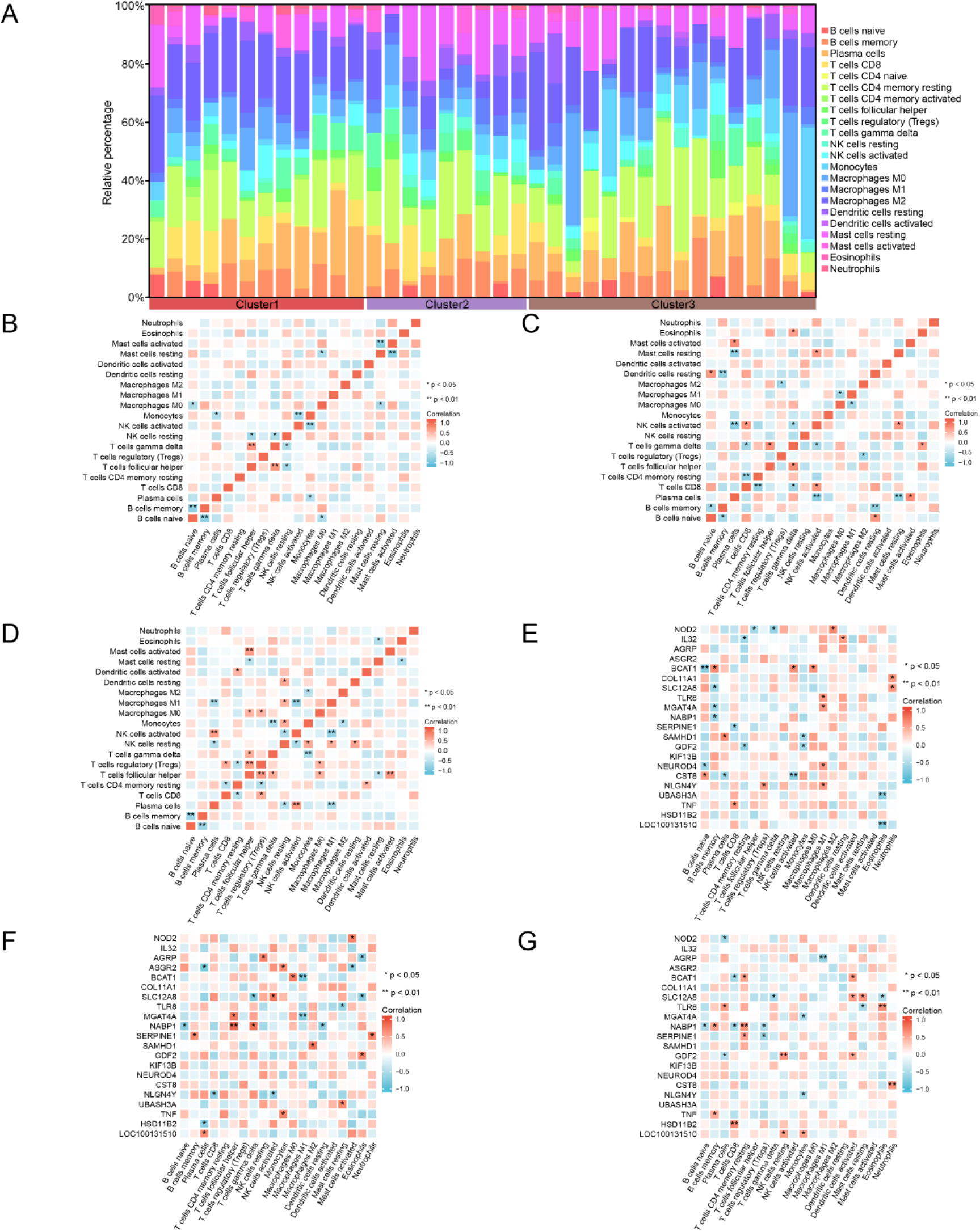
Consensus Clustering Immune Infiltration Analysis by CIBERSORT Algorithm A. Histogram of the percentage of immune cells. B-D. Correlation analysis of the abundance of immune cell infiltration in Endometriosis Cluster 1 (B), Cluster 2 (C) and Cluster 3 (D). E-G. Correlation of the abundance of immune cell infiltration in Endometriosis Cluster 1 (E), Cluster 2 (F) and Cluster 3 (G). Heat map of the correlation between the abundance of immune cell infiltration and KMGs. Cluster 1 in red, Cluster 2 in purple and Cluster 3 in brown.

Finally, the relationship between KMG expression and immune cell infiltration levels within each endometriosis subtype was investigated using correlation heatmaps (Fig.18E-G). In Cluster 1 (Fig.18E), M1 macrophages demonstrated the strongest positive correlation with KMGs. In Cluster 2 (Fig.18F), T follicular helper cells exhibited the strongest positive correlation with NABP1. In Cluster 3 (Fig.18G), M1 macrophages demonstrated the strongest negative correlation with AGRP.

## Discussion

Endometriosis is a chronic inflammatory condition that affects women of reproductive age. It is characterized by the ectopic growth of endometrial tissue, which can lead to significant morbidity, including pelvic pain and infertility(Horne & Missmer, 2022; Li et al., 2023). The underlying causes of endometriosis are still not completely clear, though recent studies have implicated various molecular pathways, including inflammatory responses(Feng et al., 2023a; Rahmioglu et al., 2023) and cellular processes such as pyroptosis(An et al., 2024; Feng et al., 2023b). The current study provides a comprehensive bioinformatics analysis aimed at identifying Pyroptosis-Related genes (PRGs) associated with endometriosis, thereby offering novel insights into the disease’s molecular underpinnings.

Pyroptosis, an inflammatory form of programmed cell death, has been increasingly recognized as a critical player in various diseases, including cancer and autoimmune disorders(Qiao et al., 2024; Rao et al., 2022; Wang et al., 2024; Zhang et al., 2022). In endometriosis, chronic inflammation is a hallmark, and the involvement of pyroptosis in this process suggests a potential link between cellular death mechanisms and the perpetuation of inflammatory cycles within ectopic endometrial tissues(Nishimoto-Kakiuchi et al., 2023; Taylor et al., 2021). The findings from this study, particularly the identification of 26 PRDEGs in endometriosis, underscore the relevance of pyroptosis in the disease’s pathophysiology.

Previous research has highlighted the role of inflammation in endometriosis, with cytokines such as IL-1β and IL-18 being key mediators(Rahmawati et al., 2023; Yu et al., 2023). These cytokines are also crucial in pyroptosis, suggesting that the inflammatory environment of endometriosis may be conducive to pyroptotic cell death. The identification of PRDEGs like VCAM1 and LY96, which were found to have high accuracy in distinguishing endometriosis from normal tissues, aligns with existing knowledge that these genes are involved in immune cell adhesion and activation, processes integral to both inflammation and pyroptosis(Gao et al., 2023; Nie et al., 2022).

Furthermore, the study’s enrichment analysis revealing the involvement of pathways like NF-κB and IL-17 in endometriosis adds another layer of complexity to the disease’s inflammatory milieu. NF-κB, a well-known regulator of immune responses(Liu et al., 2017), is also a key player in the induction of pyroptosis(Liu et al., 2023; Luo et al., 2022), suggesting that therapeutic strategies targeting this pathway could modulate both inflammation and cell death in endometriosis.

One of the significant contributions of this study is the construction of a diagnostic model based on the expression of PRDEGs. This model demonstrated high accuracy and discriminative power in identifying endometriosis, as validated in an independent dataset(GSE25628). We classified EMs into three subtypes based on 21 KMGs through WGCNA, and identified the relationship between KMGs and immune cell infiltration in different subtypes of endometriosis. This demonstrates PRGs and immune cell infiltration play a crucial role in the occurrence and development of EMs.

This study provides valuable insights, but its reliance on publicly available data introduces potential limitations regarding sample representativeness and data integrity. Future research should prioritize validating these findings using larger, more diverse cohorts to ensure broader applicability. Secondly, the study’s bioinformatics approach, while powerful, cannot establish causality. The identification of PRDEGs and their associated pathways provides a foundation for hypothesis generation, but experimental studies are required to elucidate the exact mechanisms by which pyroptosis-related genes contributes to endometriosis.

For instance, in vivo models of Endometriosis could be used to assess the impact of modulating PRGs on disease progression and symptoms. Lastly, the therapeutic implications of targeting pyroptosis in endometriosis need to be explored. While the study suggests potential targets, such as the NF-κB pathway, the safety and efficacy of such interventions remain unknown. Given the complexity of endometriosis, a multifaceted approach that combines pyroptosis modulation with existing anti-inflammatory and hormonal treatments may be necessary.

## Conclusions

This study provides novel insights into the role of pyroptosis in the pathogenesis of endometriosis. We identified key pyroptosis-related differentially expressed genes (PRDEGs) and developed a diagnostic model with potential clinical utility. These findings warrant further investigation to validate their robustness and generalizability across diverse patient populations. Future research should also focus on exploring the therapeutic potential of targeting pyroptosis-related pathways and molecules in endometriosis, which may lead to the development of novel and effective treatment strategies for this debilitating condition.

## Declarations

- Ethics approval and consent to participate Not applicable.
- Consent for publication Not applicable.
- Availability of data and materials https://www.ncbi.nlm.nih.gov/geo/query/acc.cgi?acc=GSE7305 (GSE7307 and GSE11691)
- Competing interests The authors declare that they have no competing interests.

- Funding This work was supported by the grants from National Natural Science Foundation of China (No. 82201820) and Scientific Research Fund for Introduced Talents of the First Affiliated Hospital of Wannan Medical College (No.YR20220220).

- Authors’ contributions These should be presented as follows: PT, YL and JD designed the research study. PT and LW performed the research. KG, XL and CS analyzed the data. CD and GN reviewed entire manuscript. PT and JD wrote the manuscript. All authors contributed to editorial changes in the manuscript. All authors read and approved the final manuscript.

## Acknowledgements

Not applicable.

